# Attention defines the context for implicit sensorimotor adaptation

**DOI:** 10.1101/2024.09.03.611108

**Authors:** Tianhe Wang, Jialin Li, Richard B. Ivry

**Author notes:** Corresponding author: Tianhe Wang.

## Abstract

Movement errors are used to continuously recalibrate the sensorimotor map, a process known as sensorimotor adaptation. Here we examined how attention influences this automatic and obligatory learning process. Focusing first on spatial attention, we compared conditions in which the visual feedback that provided information about the movement outcome was either attended or unattended. Surprisingly, this manipulation had no effect on the rate of adaptation. We next used a dual-task methodology to examine the influence of attentional resources on adaptation. Here, again, we found no effect of attention, with the rate of adaptation similar under focused or divided attention conditions. Interestingly, we found that attention modulates adaptation in an indirect manner: Attended stimuli serve as cues that define the context for learning. The rate of adaptation was significantly attenuated when the attended stimulus changed from the end of one trial to the start of the next trial. In contrast, similar changes to unattended stimuli had no impact on adaptation. Together, these results suggest that visual attention defines the cues that establish the context for sensorimotor learning.

## Introduction

Adapting our movements in response to changes in the environment and body is essential for survival. A deer escaping a predator needs to adjust the force used to jump over a fence depending on the resistance of the underlying surface. To maintain stability, a bird will adjust how fast to flap its wings as a function of the prevailing winds (Pennycuick, 1996). Sensorimotor adaptation involves multiple learning systems, including an implicit system that operates to maintain the calibration of the sensorimotor map and an explicit system that can be flexibly deployed to deploy strategies that minimize performance error. (J. A. Taylor et al., 2014; Jordan A. Taylor & Ivry, 2014). While explicit mechanisms are critical for the volitional changes in behavior, the implicit processes keep the motor execution system precisely calibrated in an automatic way.

Prior studies have shown that sensorimotor learning systems are sensitive to attentional manipulations. For example, the rate of adaptation is reduced when participants are engaged in a concurrent, secondary task. This phenomenon has been observed in all kinds of sensorimotor adaptation tasks including prism adaptation (Redding & Wallace, 1996), visuomotor rotation (T. S. L. Wang et al., 2022), and force-field adaptation (Jordan A. Taylor & Thoroughman, 2007; Thoroughman et al., 2007). The slower rate of adaptation observed under dual-task conditions is usually attributed to a limitation within the explicit system (Song, 2019; Jordan A. Taylor & Ivry, 2014). For example, the cognitive resources required to complete the secondary task limit that which is available for discovering a strategy to counteract the perturbation.

The implicit recalibration process, on the other hand, is assumed to operate in an automatic manner. It uses a forward model to predict the sensory consequences of a movement and the sensory prediction error, the difference between the predicted and actual feedback, to recalibrate the system. Implicit adaptation has been demonstrated to be a remarkably robust process. It is observed even in situations in which there is no explicit task-error or when participants are instructed to disregard the feedback and move directly to the target (H. E. Kim et al., 12/2018; Mazzoni & Krakauer, 2006; Morehead et al., 2017).

However, it is less clear that whether the implicit adaptation system is influenced by attention. While we tend to assume that automatic processes do not utilize cognitive resources, this proposal has never been systematically explored in the domain of implicit sensorimotor adaptation. There is a substantial body of evidence showing that implicit processes can be susceptible to the effects of divided attention. For instance, visual priming is diminished when participants are engaged in a secondary task, an effect attributed to a reduction in the depth of processing of the priming stimuli (Otsuka & Kawaguchi, 2007; Prull et al., 2016; Schmitter-Edgecombe, 1996). Divided attention might impact implicit sensorimotor adaptation in a similar way.

Beyond being subject to limits on available cognitive resources, there are other ways in which attention could influence implicit adaptation. For instance, the focus of visuospatial attention could have a modulatory effect on the processing of a visual feedback signal. This hypothesis is motivated by studies in which multiple feedback cursors are simultaneously presented during sensorimotor adaptation. Under these conditions, the functionally effective error signal is a composite signal, one that corresponds to the averaged error conveyed by the feedback (Kasuga et al., 2013). However, if the participant is instructed that one of the feedback cursors is more task-relevant, the implicit system allocates a larger weight to this input when calculating the composite error (J. Tsay et al., 2024).

A third way by which attention might influence implicit adaptation is through context mediation. Motor memories are known, to some degree, to be context specific, with the manifestation of the memory reduced when the context changes (Heald et al., 2021, 2022; Shea & Morgan, 1979). For example, there is limited generalization of the adapted behavior when the target changes shape (Poh et al., 2021) or when participants switch from reaching with their arm to reaching with a tool (Kluzik et al., 2008). We hypothesize that attention might influence how the context is established: Attended stimuli may serve as cues that define the context for learning. Thus, the manipulation of attention might result in a contextual change, with a consequence being reduced expression of adaptation.

In the current study, we examine how attention influences implicit sensorimotor adaptation, focusing on the three hypotheses mentioned above. The primary task was always a center-out reaching task in which we used a perturbation method that restricts learning to implicit adaptation. To manipulate attention, we used a secondary task in which participants were required to make a perceptual discrimination concurrent with the reaching task. To examine the effect of visuospatial attention, the stimuli for the discrimination task were placed either near or far from the feedback cursor. To examine the effect of cognitive resources, we varied the difficult of the secondary task. Finally, to examine the hypothesis that attention impacts adaptation by re-defining the learning context, we manipulate the consistency of attended stimuli across trials. Taken together, these experiments provide a comprehensive picture of how implicit adaptation is influenced by attention.

## Results

To isolate implicit adaptation, we used task invariant, clamped feedback during a visuomotor adaptation task (Morehead et al., 2017; J. S. Tsay, Ivry, et al., 2021). Using a web-based platform and their personal computer system, participants made center-out movements by moving their finger across a trackpad (Fig 1a). Feedback was conveyed by the position of a visual cursor, presented for a brief interval when the amplitude of the movement reached the target distance. After a familiarization block in which the position of the cursor was aligned with the movement, we perturbed the feedback. On these trials, the cursor was displaced by a pre-determined angle relative to the target; thus, the angular position of the feedback was not dependent on the position of the participant’s hand. The participant was fully informed of the nature of the feedback and instructed to ignore it. Nonetheless, the “sensory prediction error” introduced by this clamped feedback causes the next movement to be shifted in the opposite direction (Avraham et al., 2021, 2022; H. E. Kim et al., 12/2018; Morehead et al., 2017; J. S. Tsay et al., 2020). This change occurs automatically and outside awareness, with all the hallmarks of pure implicit adaptation.

**Fig 1.**
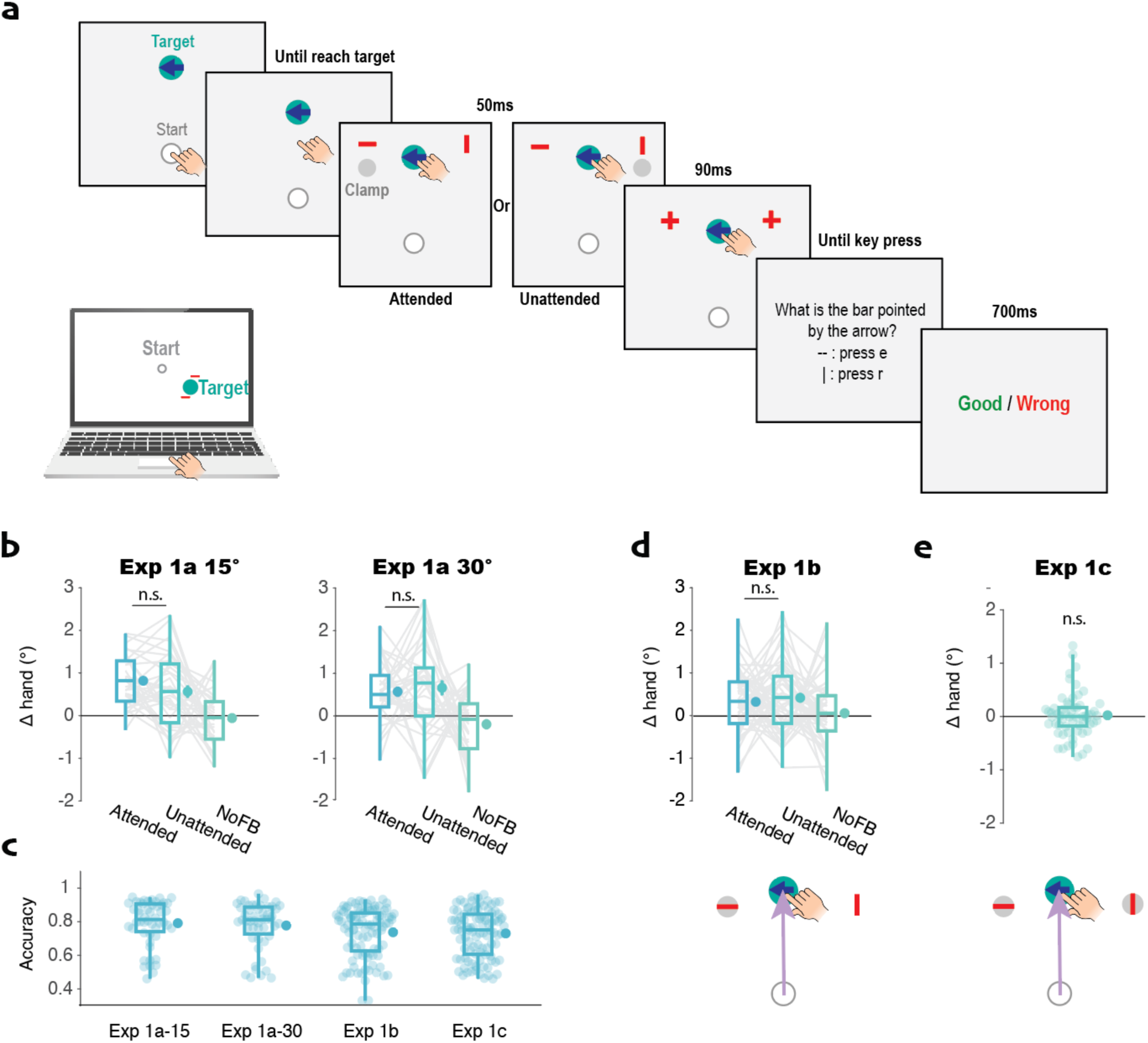
Trial-by-trial implicit adaptation is not modulated by visual spatial attention. a) Event sequence of Experiment 1a: Participants made a reaching movement towards the target. The arrow on the target indicated the trial-relevant bar for the orientation discrimination task. The bar and clamped feedback cursor were presented simultaneously when the radial distance of the hand movement reached the target. After 50 ms, the bars were converted to “+” signs, serving as pattern masks. Following a keyboard entry to indicate the orientation of the cued bar, feedback was provided. b) Δhand angle in Experiment 1a. The presence of either a 15° (left) or 30° (right) clamped cursor induced implicit adaptation. The magnitude was similar for the attended and unattended conditions. c) Accuracy on the orientation discrimination task for Experiment 1. d) Δhand angle (top row) and example of stimulus display showing bar embedded in clamped cursor (bottom row) for Experiment 1b. Again, adaptation was not influenced by the focus of spatial attention. e) Adaptation in response to the cursor on the cued side was abolished when a second clamped cursor was presented on the opposite side in Experiment 1c. For the boxplot, whiskers extend to ± standard deviation. The dot on the right of the box represents the mean and the bar represents the standard error. This format is used in all of the figures.

To manipulate visuospatial attention in Experiment 1, we had participants perform a visual discrimination task with the reaching task. When the amplitude of the hand movement reached the target distance, two bars appeared simultaneously with the clamped feedback (Fig 1a). One bar appeared just above the clamped feedback; the other appeared on the other side of the target. The orientation of each bar was randomly and independently set to be either vertical or horizontal, and the bars were masked after 50 ms by turning into identical crosses. The participant was required to report the orientation of the cued bar. The cue was an arrow that was positioned within the target and visible from the start of the trial. Since the bars were small (5 pixels long) and briefly presented (50 ms), we assumed that participants would shift their attention (and eyes) to the cued location. Mean accuracy on the secondary task was 79.2% in Experiment 1, indicating that the secondary task was demanding (Fig 1c, Fig S1). The analyses were limited to participants with accuracy >=70% on the visual discrimination task, a criterion designed to select only those who followed the attentional manipulation (see Table S1 for the exclusion percentage). With this design, we asked if the magnitude of adaptation differed as a function of whether the clamped feedback appeared adjacent to the cued bar or distant from the cued bar.

Visuospatial attention does not influence implicit adaptation.

In Experiment 1a, the clamped feedback was presented 15° from the target, with the direction (clockwise or counterclockwise) randomized from trial to trial. As our assay of implicit adaptation, we measured the trial-by-trial change of hand angle (Δhand). Participants showed significant adaptation in response to the clamped feedback: Overall, the mean change in heading angle was of 0.77° ± 0.12° (mean ± SE) on trials in which the clamp appeared near the attended bar and 0.56° ± 0.16° on trials in which the clamp was opposite the attended bar (Fig 1b). Importantly, the magnitude of adaptation did not differ between the two conditions (*t*(34)=1.2, *p*=0.24, *d*=0.20, *bf10*=0.35), suggesting that this process is not modulated by spatial attention. There was no consistent change in hand angle after the no-feedback trials, confirming that the change in hand angle after feedback trials is due to the clamped feedback and not the presence of a secondary task.

Given that we obtained a null result, we repeated the experiment, using a 30° clamp and shifted the position of the bars in a similar manner. We assumed that these more eccentric locations would make it even more incumbent for the participants to re-orient attention towards the cued location, and thus enhance any potential subtle effects of spatial attention on implicit adaptation. However, the results were essentially identical to that observed with the 15°. There was no difference in the change in hand angle between the attended and unattended conditions (*t*(37)<0.001, *p*=1.00, *d*<0.001, *bf10*<0.001, Fig 1b).

One concern with our attentional manipulation in Experiment 1a is that attention is directed to an object (i.e., the bar), and this might preclude other objects (i.e., the feedback cursor) from being part of the focus of attention, even if positioned close to the attended object (Logan, 1996; Roelfsema et al., 1998). To address this concern, we shifted the position of the red bars in Experiment 1b such that the bar and clamped cursor were overlapping on one side. As in Experiment 1a, we again failed to find a difference between the attended and unattended conditions (*t*(56)=0.11, *p*=0.91, *d*=0.06, *bf10*=0.14, Fig 1d), providing further evidence that spatial attention does not modulate implicit adaptation.

In Experiment 1c, we presented two feedback cursors, symmetrically arranged around the target, and displayed a bar within each cursor. As with Experiment 1a and 1b, an arrow embedded in the target indicated which bar was relevant for the discrimination task. We reasoned that if spatial attention influences implicit adaptation, a change in hand angle on trial n+1 would be associated with the side of the clamp/bar on trial n. In contrast to this hypothesis, we failed to find evidence of adaptation in Experiment 1c, with an average Δhand at approximately 0° (*t*(59)=0.72, *p*=0.47, *d*=0.09, *bf10*=0.18, Fig 1e). Thus, we assume the system assigned equal weight to the two symmetric cursors at ±15° which resulted in a net error of 0° (Kasuga et al., 2013).

The trial-by-trial measure of adaptation used in Experiment 1 yielded a consistent change in hand angle, but one that was small in magnitude (see Figs 1b-d). This might reduce our sensitivity to detect an attentional effect. To address this concern, we applied a clamp in Experiment 2a that appeared on one side of the target for 80% of the trials and on the other side for 20% of the trials. With this method, the effect of adaptation will accumulate across trials, with the hand direction biased to move in the opposite direction of the 80% clamp. Each participant completed two blocks of trials, one in which they were more likely to attend to the side that has the high probability of displaying the clamp (attended block) and one in which they are more likely to pay attention to the opposite direction (unattended block, see Methods for details).

We observed robust accumulated adaptation in both the attended and unattended conditions, with functions that reached an asymptote of around 10° (Fig 2b). Consistent with the results from the trial-by-trial experiments, we did not observe any difference between the attended and unattended conditions. This null effect was seen in a cluster-based permutation test comparing each point in the learning functions and when we averaged across the whole learning function (*t*(63)=0.39, *p*=0.70, *d*=0.05, *bf10*=0.15). We conducted another experiment (Experiment 2b) in which clamped feedback was presented on both sides of the target, comparing a session in which attention was cued to one direction on 80% of the trials to a session in which the cued direction was randomized across trials (50%/side). Similar to what we observed in the single-trial analysis of Experiment 1c, we did not observe any adaptation in this condition and no difference between the two attention conditions (*t*(44)=-0.35, *p*=0.73, *d*=-0.11, *bf10*=0.17, Fig 2b).

**Fig 2.**
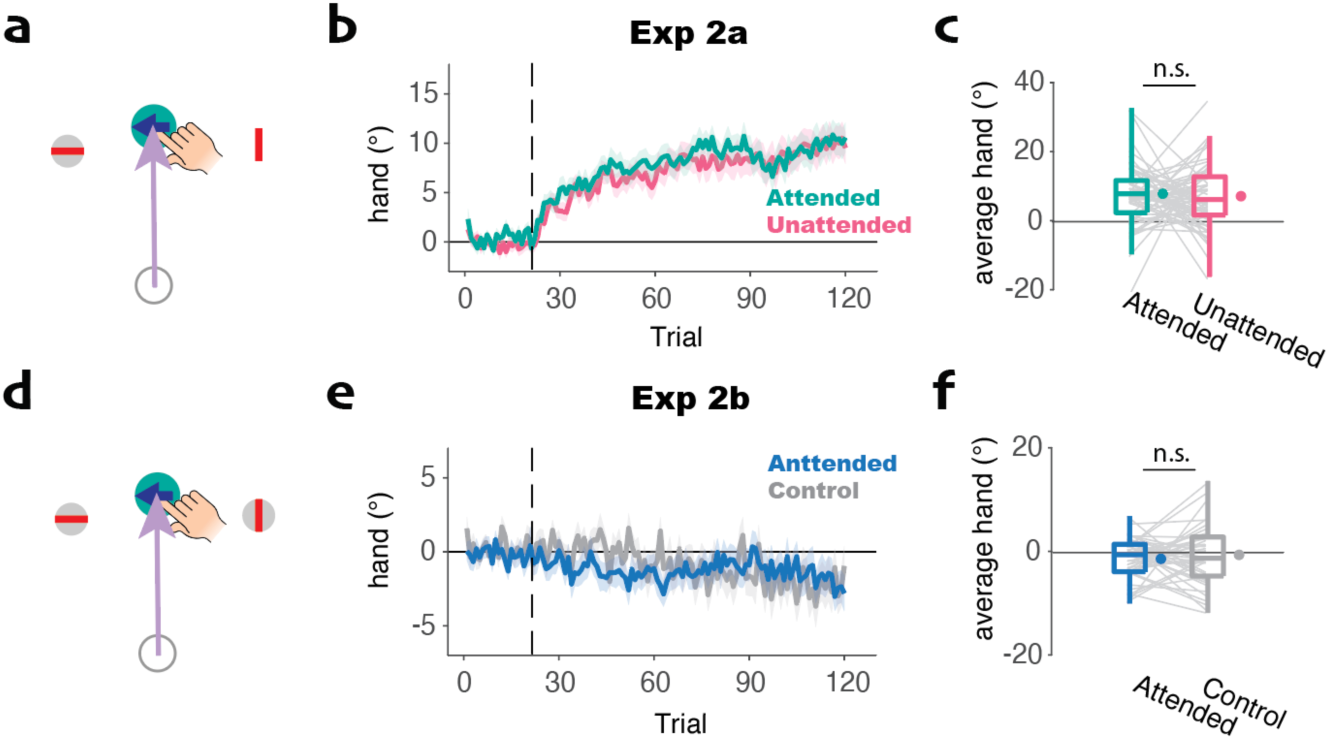
Accumulated adaptation is not modulated by visuospatial attention. a) Representative display for the attended condition Experiment 2a. The clamp appeared on the cued side on 80% of the trials and on the opposite side for the other 20%. These percentages were reversed for the unattended condition. b) Learning curve with the vertical dashed line indicating clamp onset. We applied a cluster-based permutation test to compare the learning curves; However, no significant cluster was found. c) Average hand angle across trials in which the clamp was presented. No significant difference was observed in the average hand angle. d-f) Similar to a-c but for Experiment 2b. Two clamped cursors were presented with the arrow cuing the target on one side on 80% of trials (Experimental condition). In the control condition, the direction of the arrow was randomized (50% each direction). Neither condition resulted in significant adaptation.

In summary, the results from Experiment 1 and 2 consistently show no difference in the magnitude of adaptation resulting from a feedback cursor on the attended side of space compared to when the feedback cursor is on the unattended side of space. As such, we infer that adaptation is not modulated by visuospatial attention.

### Implicit adaptation does not tax cognitive resources

Despite the absence of a modulatory effect of visuospatial attention on adaptation in Experiment 1 and 2, the size of the trial-by-trial change in hand angle in Experiment 1 is small (means around 0.5°) compared to that observed in previous experiments using the same platform but without a secondary task (means around 1.5-2°-- see (J. S. Tsay, Ivry, et al., 2021; T. Wang & Ivry, 2023; T. Wang et al., 2023). This suggests that the inclusion of a secondary task may attenuate adaptation. Alternatively, the attenuation might be due to the introduction of the additional visual stimuli (arrow and bars), inputs that could impact processing of the feedback cursor.

To compare these two hypotheses, we used a simplified secondary task in Experiment 3, one in which the display was limited to the target and clamped feedback cursor. For the dual-task condition, the participant was instructed to pay attention to the clamped feedback and report the cursor’s location relative to the target (left or right) at the end of each trial. In the control, single-task condition, the participant was told to ignore the clamp and no report was required although the participant was required to press a specific key after each movement to match the trial interval (Fig S2). In both conditions, the participants were informed that the position of the clamped feedback was not linked to their motor performance and that they should always reach directly to the target.

*A priori*, one might expect that attending to the feedback cursor in the dual-task condition would increase adaptation. However, the opposite was found: Adaptation was significantly attenuated in the dual-task condition compared to the single-task condition (*t*(59)=6.2, *p*<0.001, *d*=1.6, *bf10*=1.8*10^5^, Fig 3a). For the latter, the trial-by-trial change in hand angle was approximately 1.6°, a value similar to that observed in prior studies (J. S. Tsay, Ivry, et al., 2021; T. Wang & Ivry, 2023). In contrast, the magnitude of adaptation dropped to around 0.5° in the dual-task condition. This result further confirms that paying attention to the feedback does not boost adaption. Indeed, these results show that the inclusion of a secondary task can attenuate implicit adaptation.

**Fig 3.**
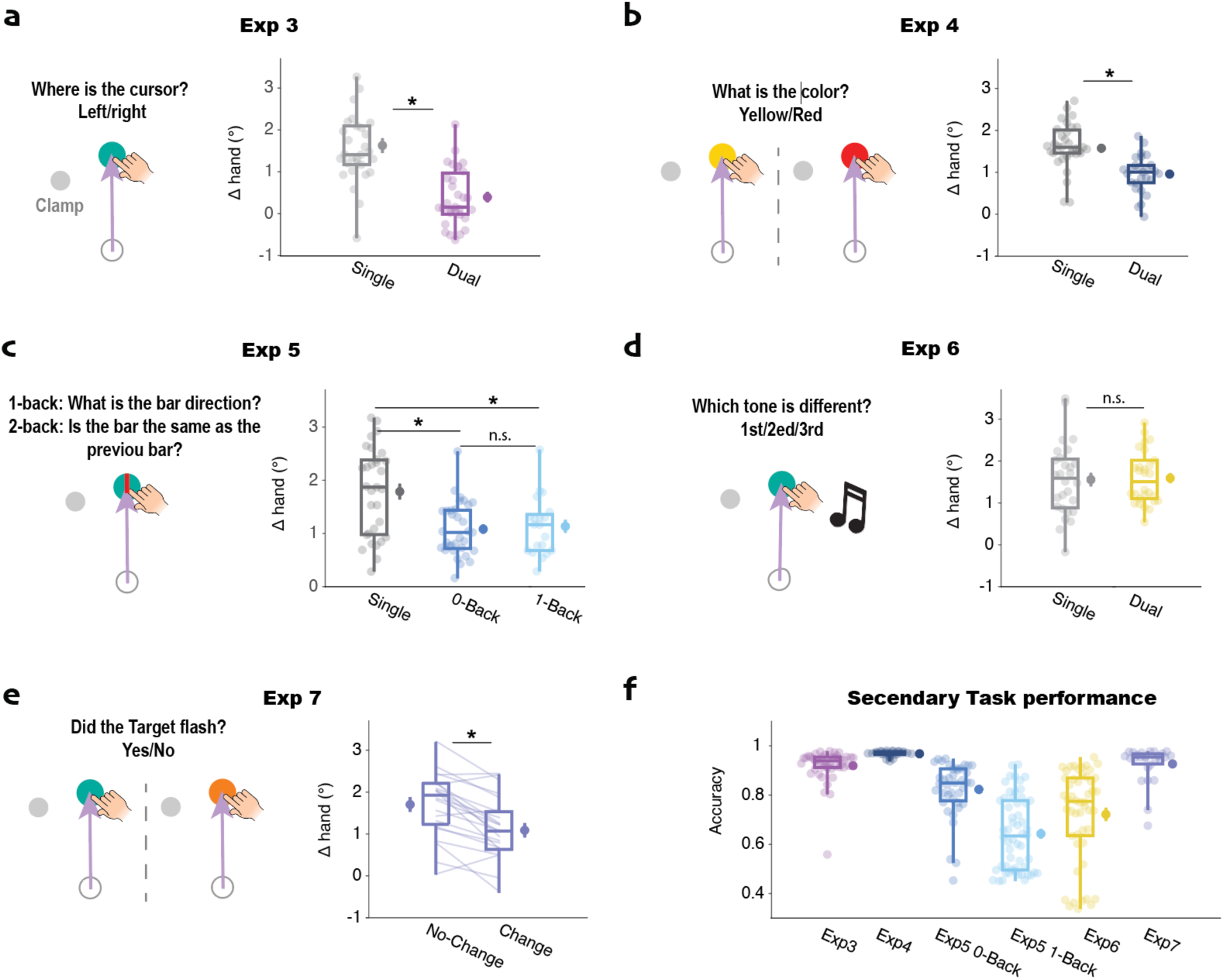
Implicit adaptation is not modulated by the availability of cognitive resources. a) Experiment 3: Adaptation was attenuated in a dual-task condition in which participants had to identify the position of the clamped cursor after completing the reach. b) Experiment 4: Adaptation was attenuated in a dual-task condition in which participants judged the color of target after completing the reach. c) Experiment 5: The bar was presented on the target and participants either ignored the bar (single-task), reported the bar orientation on the current trial (0-back, low demand dual-task), or reported if the bar orientation matched that observed on the previous trial (1-back, high demand dual-task). Adaptation was highest in the single-task condition, but the degree of attenuation did not differ between the Low and High demand versions of the dual-task conditions. d) Experiment 6: An auditory pitch discrimination task was used for the secondary task with performance compared to a group instructed to ignore the tones. Adaptation was not attenuated in the dual-task condition. e) Experiment 7: Participants reported if the target color remained constant or briefly changed to red. Although the secondary task set was similar in conditions, adaptation was attenuated on change trials compared to no-change trials. f) Accuracy on the secondary tasks of Experiments 3-7. * p<.001., n.s., not significant.

We note that in Experiment 3, the focus of attention is likely to differ between the dual and single-task conditions. In the former, the participant likely shifts attention from the target to the peripheral cursor given that they must report the location of the cursor. In the latter, the task instructions would emphasize the importance of remaining focused on the target. Similarly, the dual-task conditions in Experiments 1 and 2 also required a shift in the focus of attention from the target to a more eccentric stimulus. Perhaps the attenuation of adaptation in the dual-task conditions arises because the focus of attention (and likely fixation) changes.

To assess this hypothesis, we modified the secondary task in Experiment 4. Here the color of the target briefly changed to either yellow or red at the onset of the clamp. Participants in the dual-task condition were instructed to report the color after each movement whereas participants in the single-task were instructed to ignore the color change. Again, we observed attenuated adaptation in the dual-task condition compared to the single-task condition (*t*(59)=4.5, *p*<0.001, *d*=1.1, *bf10*=510, Fig 3a). Thus, the reduced efficacy of adaptation does not appear to be due to a dual-task-induced change in the location of attention.

Why might a secondary task attenuate adaptation? One possibility is that the attenuation might be the result of resource sharing between the primary reaching task and the secondary task. It is generally assumed that implicit adaptation does not tax cognitive resources given that the response is automatic and, indeed, cannot be suppressed even when the feedback is task-irrelevant (e.g., clamped) or the response is counter-productive to task performance (Mazzoni & Krakauer, 2006; Morehead et al., 2017). However, even if implicit and obligatory, it is still possible that adaptation draws on cognitive resources. If so, the efficacy of this process might be reduced when these resources are shared with a concurrent task.

To test this hypothesis, we performed three experiments. In Experiment 5, a red bar, oriented vertically or horizontally, appeared on the target, concurrent with the clamped cursor. We compared three conditions in which we varied the processing demands associated with the secondary task. In the control condition, participants were instructed to ignore the red bar (single-task condition). In the low demand condition, the participant indicated the orientation of the bar right after a movement (0-back). In the high demand condition, the participant reported if the orientation of the bar matched that shown on the previous trial (1-back task). The accuracy data confirmed our expectation that the 1-back condition is more difficult, and presumably, more demanding on cognitive resources: Accuracy on the orientation task dropped to 67% in the high demand condition compared to the 86% in the low demand condition (Mann-Whitney U test, *z*=5.4; *p*<0.001).

As seen in our previous experiments, adaptation was lower in both of the dual-task conditions compared to the single-task condition (1-back: *t*(49)=3.2, *p*=0.002, *d*=0.93, *bf10*=16.7; 0-back: *t*(57)=3.7, *p*<0.001, *d*=0.98, *bf10*=65.6); thus, the inclusion of a secondary task attenuated adaptation. However, the level of adaptation was similar for the high and low conditions despite the difference in difficulty (Fig 3b; *t*(42)=0.09, *p*=0.93, *d*=0.08, *bf10*=0.29). Thus, the degree of attenuation does not seem to be related to the resource demands of the secondary task, at odds with the hypothesis that adaptation is modulated by the availability of cognitive resources.

We next asked if the attenuating effect of a secondary task would also be observed if that task involved a different modality. In Experiment 6, we used an auditory secondary task, presenting at movement onset a sequence of three 40 ms tones, with the tones separated by an interval of 50 ms. Two tones were at one pitch and the third was an octave higher. Participants in the dual-task condition indicated the ordinal position of the oddball pitch whereas participants in the single-task condition were instructed to ignore the auditory stimuli. Surprisingly, the magnitude of adaptation was comparable in the two conditions (*t*(55)=0.31, *p*=0.75, *d*=0.08, *bf10*=0.28, Fig 3c) and the magnitude of adaptation in both conditions was comparable to that observed in in the single-task conditions of Experiment 3 and 4. These results also argue against a general resource account.

In the third test of the resource hypothesis (Experiment 7), we used a within-participant design, keeping the attentional demands constant while manipulating the visual display. We used a detection task in which, at the onset of the cursor feedback, the color of the target briefly changed from blue to red on 50% of the trials and remained blue on the other trials. After completing the reaching movement, the participant indicated if they had detected a color change. Given that the attentional set should be similar on change and no-change trials, we would expect a similar adaptation response if the magnitude of adaptation is influenced by cognitive resources. However, we observed reduced adaptation following change trials compared to no-change trials (*t*(21)=5.3, *p*<0.001, *d*=1.26, *bf10*=867, Fig 3e). Indeed, adaptation following the no-change trials was similar to that observed in the previous single-task conditions.

Considered as a package, the results of Experiments 5-7 fail to support the hypothesis that implicit adaptation is modulated by the availability of cognitive resources. The magnitude of adaptation was not influenced by the difficulty of a secondary task, nor did we observe attenuation of adaptation when the secondary task was in a different modality. As a post-hoc analysis, we compared all the relevant conditions from this report (see Figure S3). This analysis showed no correlation between the difficulty of a secondary task and the rate of adaptation, providing further evidence that adaptation does not require cognitive resource.

### Attention influences adaptation by defining the context

In reviewing the conditions that produced attenuation of implicit adaptation, we recognized one consistent feature: The stimulus that was the focus of visual attention at the end of the trial n-1 is different compared to the stimulus that is the focus of visual attention at the start of trial n (Fig 4d). For example, each dual-task trial in Experiment 1 ends with a report of the orientation of a bar whereas the bars are not part of the display at the start of the next trial. Another, more compelling example comes from Experiment 3. Although the displays are identical on the dual- and single-task trials, the focus of attention shifts to the clamped cursor in the former whereas it (at least as instructed) remains focused on the target in the latter. Whereas we find attenuation when there is a shift in the focus of attention, no attenuation is observed when the focus of visual attention is unchanged across trials. This distinction is apparent when considering the conditions of Experiment 7. Attenuation is only observed when the target color had changed on trial n-1.

**Fig 4.**
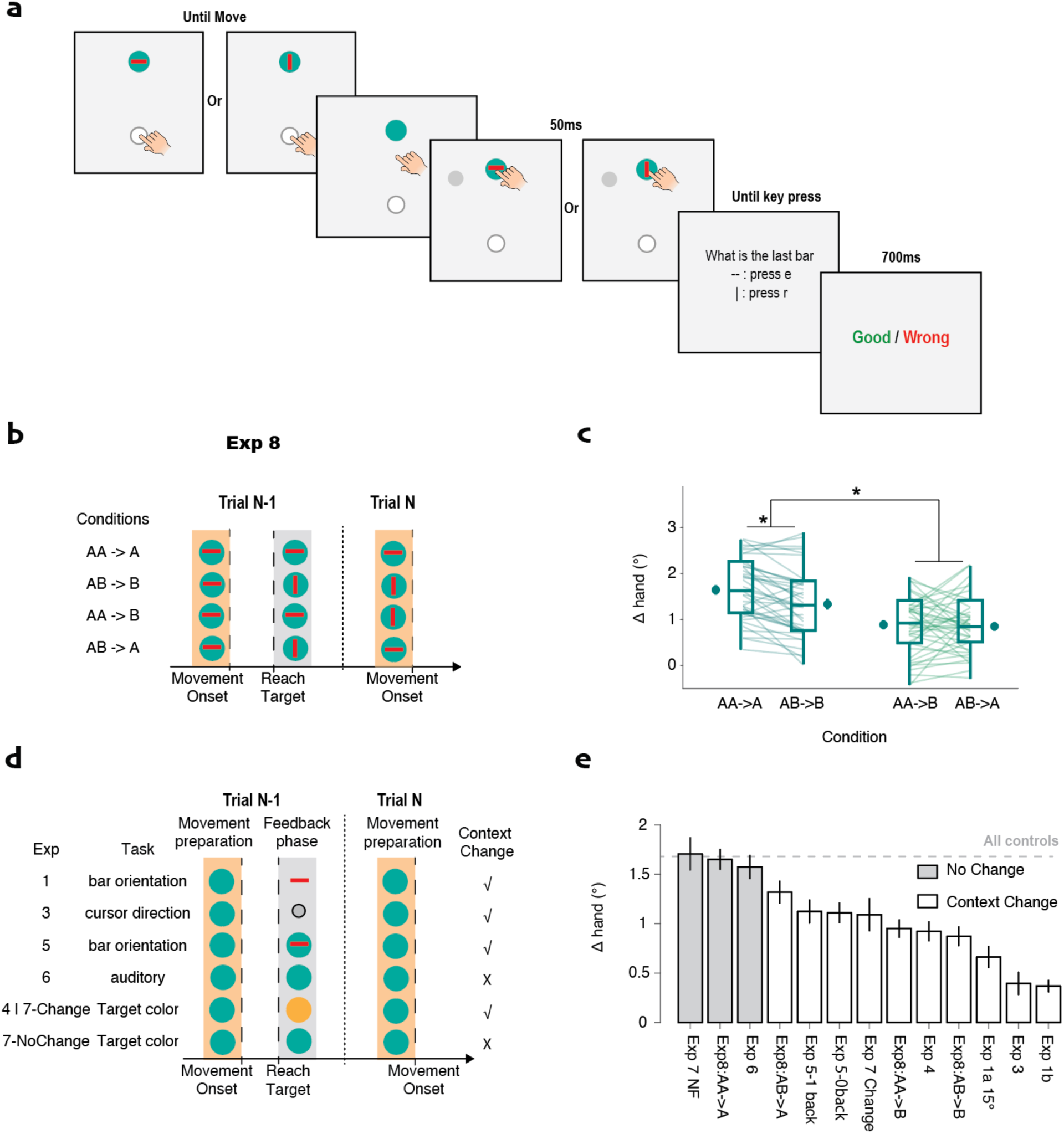
Implicit adaptation is attenuated when the attended stimulus changes between the feedback phase of trial n-1 and movement planning phase of trial n. a) Experiment 8: A bar, oriented horizontally or vertically, is presented on the target at trial onset (movement preparation) and disappears at movement onset. A second bar appears with the feedback cursor, either in the same or different orientation. The participant reports the orientation of the second bar. b) Truth table showing how bar orientation at different phases determines whether context is consistent or changes. c) The no-change (AA->A) condition shows larger adaptation compared to the other three conditions. d) Summary of contextual change in the relevant dual-task conditions of Experiment 1-7. e) Δhand, rank ordered by magnitude of adaptation with shading to indicate conditions in which the context is unchanged. “All controls” refers to the average of the single-task conditions. Error bars indicate standard error.

Why might this change in the focus of attention produce an attenuation of adaptation? We propose that the focus of attention defines the context for the memory. The implicit system is using the clamped feedback to update an internal model. These models are, to some degree, context specific. Context can be defined spatially; as shown in many studies of generalization, adaptation to movements on the right side of space minimally affects movements on the left side of space (John W. Krakauer et al., 2006, 2000; Morehead et al., 2017; Zhou et al., 2017). But context can also be defined by environmental features or internal states. For example, different internal models can be engaged for the same movement when non-compatible perturbations are contextually associated with distinct visual cues (Avraham et al., 2022; Howard et al., 2013).

We propose that our attentional manipulations had the effect of associating distinct contextual cues with internal models for adaptation. When the focus of visual attention changed, adaptation was attenuated because there was limited transfer between the two models/contexts. For example, an error used to update a model of the environment that features an orientated bar in trial n-1 will not fully generalize to a model of an environment in which the bar is not present at the start of trial n. Following this logic, presenting the same bar during movement preparation in trial n should mitigate the attenuation effect.

To test this hypothesis, we manipulated the consistency of the contextual cue across trials in Experiment 8 (Fig 4a). A bar, oriented vertically or horizontally, appeared on the target at the start of the trial and disappeared at movement onset. Another bar, with either the same or different orientation, appeared on the target concurrent with the clamp. After the reach, the participant reported the orientation of the second bar. With this design, we assume that the participants attended to both bars: The first one because they need to attend to the target to determine the reach direction and the second one because it is requirement of the secondary task.

When considered as contextual cues, the first bar defines the context for memory expression, while the second bar defines the context for memory updating (Fig 4b). If the orientation of the second bar (during feedback) on trial n-1 is the same as the orientation of the bar at target onset on trial n (AA->A and AB->B), the retrieved internal model is the same as the one that was just updated. In contrast, when the orientation at target onset on trial n differs from that at feedback on trial n-1 (AA->B and AB->A), the context has changed. Thus, the retrieved internal model is not identical to the one that was just updated. As such, we would expect to observe reduced adaptation due to limited generalization (Howard et al., 2013, 2015).

The results of Experiment 8 strongly support this idea. A mixed-linear model revealed that adaptation is significantly reduced when the bars change orientation at the onset of trial n (t(159)=4.9, p<.001, Fig 4c). Post-hoc t-tests showed that adaptation in both the AA->A and AB->B conditions was greater than in the AA->B and AB->A conditions, respectively (t’s>2.8, *ps*FDR<0.007, ds>0.61, *bf10s*>6.3). Interestingly, we also observed a slight decrease in adaptation in the AB->B condition compared to the AA->A condition (t(39)=4.8, *p*FDR<0.001, d=8.0, *bf10*=890). We hypothesize that this also reflects a contextual change given that the bar orientation is different at target onset on trials n-1 and n. However, there was no significant difference between the AA->B and AB->A conditions (t(39)=0.06, *p*FDR=1.0, d=0.06, *bf10*=0.23), despite the fact that the latter condition involves the same internal model at trial onset. We assume that adaptation is attenuated in AB->A condition due to the context change resulting from a change in bar orientation during memory updating on trial n−1 compared to the bar orientation for memory expression on trial n.

These results motivated us to review all of the relevant dual-task conditions across experiments in terms of the contextual change hypothesis (Fig 4e). Notably, in all conditions in which the context remains consistent, the magnitude of adaptation is similar and falls within the mean value of adaptation observed in single-task conditions. In contrast, attenuation is observed whenever there is a change in context cues. Interestingly, the dual-task conditions in Experiments 1 and 3 demonstrated the strongest attenuation. These are the two conditions in which the spatial focus of attention (and presumably fixation) changed from the end of trial n-1 to the start of trial n. Such spatial shifts may define a more substantial context change.

## Discussion

The current study addresses whether and how visual attention modulates implicit sensorimotor adaptation. Taken as a package, the results reveal three key findings. First, implicit adaptation does not tax attentional resources as variation in the difficulty of a secondary task had no influence on adaptation across the set of experiments. Second, implicit adaption was not directly modulated by the focus of visuospatial attention: The rate and extent of implicit adaptation were unaffected by whether or not participants paid attention to the feedback cursor. Third, attention can modulate adaptation in an indirect manner. Specifically, stimuli that are the focus of attention serve as cues to help define the learning context, and this context constrains how the sensorimotor map is updated and expressed. When the attended stimuli change across trials, learning is attenuated due to imperfect generalization between non-identical contexts.

### Attention decides the relevant contextual cue for sensorimotor memory

Context has long been recognized as an important constraint on learning. A classic example from described in introductory psychology textbooks is that test performance improves when an exam is given in the same room as where the class was taught, and even more so, when the student takes the same seat (Godden & Baddeley, 1975; Smith, 1984). The importance of context has been observed with all sorts of species in learning studies ranging from simple conditioning to complex skill learning. This body of work indicates that the learned content is associated with a specific context and can be more easily accessed when that context is activated(Smith & Vela, 2001).

The relevance of context-dependent learning has also been explored in studies of sensorimotor adaptation. Perhaps most dramatic are studies of showing limitations in the generalization of implicit adaptation(John W. Krakauer, 2009; John W. Krakauer et al., 2006). Initially, the limits on spatial generalization were described in terms of tuning functions (J. W. Krakauer et al., 1999; J. Wang & Sainburg, 2005); however, subsequent studies showed that generalization for movements in the same direction was also limited if the action was performed with different tools (Kluzik et al., 2008), or even if the target shape changed across trials (Poh et al., 2021). These observations motivated the idea that the sensorimotor system develops distinct internal models for different contexts, and that generalization is dependent on the degree of overlap between contexts. Thus, even when the trials are interleaved, participants can even learn to respond to perturbations in opposite directions if the movements are associated with distinct contextual cues (Howard et al., 2015).

However, the criteria that define the contextual cues for sensorimotor learning remain unclear. The seminal work of Howard and colleagues highlighted how spatial and/or dynamic cues were more important for defining the context for sensorimotor adaptation compared to simple, static cues such as color (Howard et al., 2013, 2015). Subsequent studies have shown that abstract features such as decision uncertainty can serve as contextual cues (Ogasa et al., 2024). Most relevant to the current study is the work of Song and colleagues, showing that attentional diversion can also be a context for adaptation (Im et al., 2016; Song & Bédard, 2015). In particular, generalization was greater when participants continued to do a secondary task compared to a condition in which they switched to a single task condition (Liddy & Song, 2022; Song & Bédard, 2015).

While there is a substantial empirical literature showing the features that can (and cannot) serve as contextual cues, the underlying principles determining their effectiveness remain unclear. The results from the current study help define what constitutes an effective contextual cue for implicit adaptation. Taken as a whole, the results from our eight experiments suggest that for a visual cue to effectively define a unique context, that cue must be attended to by participants either while they are preparing for the movement (memory retrieval) or receiving feedback about the movement (memory updating). Crucially, the attended cue need not be part of the motor task to define the context for adaptation. As shown in many of our experiments, adaptation was attenuated when the stimulus for the secondary discrimination task was not relevant for the reaching movement (e.g., the target or feedback cursor). Conversely, a visual cue that is unattended failed to influence adaptation, presumably because such task-irrelevant cues are not part of the context.

Our hypothesis that attention has an indirect effect on adaptation by shaping the learning context provides a fresh perspective when considering previous findings on contextual cues. A large number of studies have shown how our ability to learn interleaved perturbations varies in a fairly dramatic way as a function of the cues used to define those perturbations (Howard et al., 2013, 2015). We would argue that the varying effectiveness of different stimuli tested in these studies reflects their capacity to engage attention. Similarly, abstract properties such as cognitive demands or task uncertainty can serve as effective contextual cues because they are critical features of the secondary task and thus, a target of attention (Ogasa et al., 2024; Song & Bédard, 2015). Avraham et al. (Avraham et al., 2022) argued that an effective contextual cue must be in temporal proximity with the movement, a requirement to ensure that is bound to the motor command. However, in Avraham et al., the contextual cues also served as the imperative signal, requiring participants to attend to these cues during movement preparation, which may explain why those stimuli were validated as contextual cues.

One puzzle here is the absence of attenuation in Experiment 6 where we used auditory stimuli for the secondary task. Here we failed to observe a difference between the dual and single task conditions event though the auditory tones had to be attended to in the former. One hypothesis is that the auditory stimulus automatically captures attention (Dunifon et al., 2016), even though it is only task-relevant in the dual task condition. However, if this were so, we would have expected that implicit adaptation would have been attenuated in both conditions whereas, when compared to other experiments, the level of adaptation is similar to the single task conditions. Alternatively, it may be that the modality separation between the primary (clamp) and secondary (tone) stimuli precluded the tones from defining a context for sensorimotor adaptation. One weakness with this hypothesis comes from the work of Avraham et al.(Avraham et al., 2022). In that study, visual and auditory cues were used to define the context for adaptation. We note that in the Avraham et al. study, the auditory cue was relevant for the adaptation task; it served as the imperative. In contrast, in the present work, the auditory cue was only relevant for the secondary, perceptual discrimination task.

Another weakness with our online experiments is that we did not measure eye movements. As such, our inferences about the focus of attention are indirect and we cannot say how the observed attentional effects (or lack thereof) might be influenced by eye position. We assume that participants fixate on the target in the single-task situations, an assumption consistent with studies showing that participants look at the target during reach planning and execution (Bromberg et al., 2019). Moreover, the onset of the target served as the imperative and its disappearance indicated that participants needed to recenter their hands. As such, we assume participants shift their gaze to the target prior to the reach. Similarly, for the secondary tasks in which we manipulated the orientation of a bar, we assume that participants shift their gaze to the bar given that it is small and only shown for a short duration (50 ms). What is unclear are the fixation patterns on trials in which the secondary stimuli are unattended, which needs to be examined by future works.

### Sensorimotor adaptation is automatic and free from attentional resource

While we have shown how attention can modulate adaptation in an indirect manner by defining the learning context, our results also underscore the robustness of the implicit sensorimotor learning system. Previous research has shown that this implicit learning process operates in an obligatory manner; it cannot be suppressed even when the “error signal” is irrelevant to the task or when the operation of implicit adaptation actually degrades task performance (Mazzoni & Krakauer, 2006; Morehead et al., 2017). However, the fact that the process is obligatory does not preclude the possibility that it could be modulated by attention. For example, paying attention to the feedback might strengthen the error signal and enhance adaptation (J. Tsay et al., 2024).

Strikingly, our results indicate that implicit adaptation is unaffected by the focus of visuospatial attention. The rate and magnitude of adaptation was indistinguishable between conditions in which attention was focused in the vicinity of the feedback cursor compared to when attention was directed to the opposite side of the target, even when this distance spanned 60°. Moreover, we failed to observe an effect on implicit adaptation when the clamped cursor itself was the secondary stimulus compared to when participants were told to ignore this stimulus (Experiment 3).

The results also show that implicit adaptation does not draw on attentional resources. The learning rate was not directly influenced by inclusion of a secondary task, a result that held for tasks spanning different modalities (auditory/visual) and functions (discrimination and working memory tasks). This resilience to divided attention is striking especially when considering that many other implicit operations do appear to be sensitive to the availability of attentional resources. For example, the encoding of implicit perceptual and semantic memories typically suffers under divided attention as evidenced by reduced priming effects (Rajaram et al., 2001; Wolters & Prinsen, 1997).

Why is the implicit sensorimotor system not modulated by attention? One possibility is that the system encodes the error in an all-or-nothing manner. As such, the computation of the sensory prediction error may require only superficial processing of the visual feedback to reach maximal reliability. This idea is support by the observation that the magnitude of the error correction is insensitive to error size over a large range (10°-90°)(H. E. Kim et al., 12/2018) and signal clarity/salience as long as the direction of the error can be reliably computed (J. S. Tsay, Avraham, et al., 2021).

The cerebellum is a key structure for implicit sensorimotor adaptation(Kim et al., 2021; Raymond & Lisberger, 1998; Shadmehr & Krakauer, 2008). The robustness of implicit adaptation may also reflect some degree of modularity between the cerebellum and cortex. A separation between systems involved in action selection and movement implementation may enable the implicit adaptation to keep the sensorimotor system precisely calibrated in a resource-free manner, freeing up attention to support the more flexible components of our mental world.

## Methods

### Participants

All of the experiments were performed online. Using the website prolific.co, we recruited a total of 929 participants (age: 27.3 ± 4.9 years old; 492 female), with the only inclusion criteria being that the participant was right-handed with normal or corrected-to-normal vision based on their responses to the Prolific pre-screening questionnaire. For the analyses, we excluded participants who failed to meet our *a priori* performance requirements on the attention tasks (specified below). This resulted in a final pool of 669 participants (355 females). Table S1 provides details on the sample size for each Experiment. The participants were paid around $8/h.

### Apparatus

The experiments were run on OnPoint, an online platform we have developed for studies of sensorimotor control and learning (Tsay, Ivry, et al., 2021). OnPoint is written in JavaScript and accessed via the Google Chrome web browser. For these studies, the participant accesses OnPoint from their personal computer. Visual stimuli are presented on their monitor and the movement data are based on finger movements across their trackpad. The data are stored on Google Firebase, a cloud-hosted database.

As described below, the instructions were provided over a series of text screens. To ensure that participants read the instructions, a key code was embedded on one page and the participant was required to enter the code before they could start the task.

### Design

We used clamped feedback in a visuomotor adaptation task, a method designed to isolate learning to implicit sensorimotor adaptation (H. E. Kim et al., 12/2018; Morehead et al., 2017). We employed a variety of secondary tasks to manipulate different aspects of attention.

#### Clamp rotation task

To start each trial, the participant moved a white cursor into a white start circle (radius: 1% of the screen height) positioned in the center of the screen. After 500 ms, a blue target circle (radius: 1% of the screen height) appeared. The radial position of the target was set to 40% of the screen height and angular position of the target was fixed at -45° (where 0° corresponds to the rightward direction). We opted for the -45° position (downward to the right) because the motor bias is smallest at this location (T. Wang, Morehead, et al., 2024).

The participant was instructed to produce a rapid, out-and-back movement, attempting to intersect the target. Feedback, when presented, could take one of two forms. On veridical feedback trials, the feedback cursor was congruent with the hand position until the movement reached the target distance. At that point, the cursor was frozen for 50 ms before disappearing. On perturbation trials, we used endpoint clamped feedback. Here the feedback cursor disappeared at movement onset and only reappeared when the radial distance of the movement reached the target distance. The angular position of the feedback cursor was invariant, displaced by 15° from the target (except in the condition in Experiment 1a in which the clamp size was 30°). That is, the position of the cursor was independent of the position of the hand, appearing at the same location on all perturbation trials. The feedback cursor remained visible for 50 ms.

The direction of the clamped were randomized across trials. Movement time was defined as the interval between the hand leaving the start circle and when the radial distance of the hand reached the target distance (40% of the screen height). If movement time was >500 ms, the message ‘Too Slow’ was presented on the screen and remained visible for 500 ms. To help guide the participant back to the start location, a white cursor (radius: 0.6% of screen height) appeared when the hand was within 30% of the target distance.

Just before the start of trials with clamped feedback (see below), we provided a set of instructions to describe this atypical perturbation. The instructions stated that the angular trajectory of the cursor would no longer be linked to their movement but rather would be fixed on all trials. The participant was to always reach directly to the target, ignoring the cursor. These instructions were repeated to make sure the participant understood the nature of the error clamp. For the first 10 trials with clamped feedback, we used a large clamp size of ±30°/±90°/180° to make clear to the participant that the feedback position was not under their control. After this familiarization phase, an instruction screen asked the participant to indicate if they were aiming for the target or another location. If the participant indicated they were reaching somewhere other than towards the target, the experiment was terminated.

We applied the same trial structure for all experiments except Experiment 2 (see below). There was a total of 560 trials beginning with a 10-trial baseline block with veridical continuous feedback. The next 10 trials were used to familiarize the participants with the clamped feedback procedure. Then the secondary task was described. The remaining 540 trials were divided such that the ±15° clamped feedback cursor appeared on 2/3 of the trials and was absent on 1/3 of the trials. The conditions were counterbalanced within each set of 30 trials.

We next describe the design of the attentional manipulations for each experiment.

#### Experiment 1

A spatial cueing task was used to manipulate spatial attention. On each trial in Experiment 1a, an arrow was presented within the target, pointing to either the left or right. When the endpoint clamp feedback appeared, two red bars were displayed, one on either side of the target. The angular deviation of the two bars was 15°, such that the feedback cursor on clamp trials was positioned beneath one of the red bars (see Fig 1a). Each bar was oriented either vertically or horizontally, and the participant’s task was to indicate the orientation of the cued bar. The feedback cursor and bars were only visible for 50 ms, after which the cursor disappeared, and the bars were transformed into two crosses for 90 ms. The latter served to mask the orientation of the bars, precluding visual aftereffects that could be used for the discrimination task. After the mask disappeared, a message appeared on the screen, indicating that the participant should press the “e” key with the left hand if the cued bar was horizontal or the “r” key with the left hand if the cued bar was vertical. Feedback on the orientation discrimination task (“Good” or “Wrong”) was presented for 700 ms before the start of the next trial. We refer to the condition in which the cued bar and feedback cursor appeared at the same location as “attended” and the condition in which the cued bar was on the opposite side of the clamp cursor as “unattended”, counterbalanced during the training session. Our key experimental question is whether adaptation will be greater in the attended condition compared to the unattended condition.

In Experiment 1a, we assumed that on attended trials, the clamped feedback cursor and cued red bar would fall within the focus of spatial attention. However, these two objects appeared at distinct locations in space; it may be that the focus of attention was associated with an object (i.e., the red bar) rather than a location. To address this issue, we presented the red bar at the same radial and angular position as the feedback cursor in Experiment 1b. Thus, on attended trials, the red bar was now presented within the feedback cursor. All other aspects of the design remained the same as in Experiment 1a.

In Experiment 1c we presented two clamped cursors symmetrically positioned around the target and embedded one red bar within each cursor. The arrow embedded in the target indicated which red bar was relevant for the orientation report. With this design, one location, and thus cursor, is more task-relevant than the other. We asked if adaptation would be induced with this type of display or if the “balanced” feedback would eliminate adaptation(Kasuga et al., 2013).

#### Experiment 2

The results of Experiment 1 failed to find any effect of spatial attention on implicit sensorimotor adaptation. One concern, however, is that we used a trial-by-trial design in each experiment: With such designs, the size of adaptation is relatively small, and thus may have limited our sensitivity to detect differences between the attended and unattended sides.

To allow adaptation to accumulate across trials, we used a pseudo-block design in Experiment 2, with the clamped cursor appearing on one side on 80% of the trials (CW or CCW, counterbalanced across participants). We again presented two bars on each trial and an arrow indicated which bar was relevant for the orientation judgment. We created attended and unattended blocks by manipulating the probability of the direction of the arrow, with the probability of the clamp location and cued direction independent of each other.

In the attended condition, the direction of the arrow cued the expected side of the cursor on 80% of the trials. As such, the cursor and cued bar were on the same side on 68% of the trials, 64% of which on the high probability side and 4% on the low probability side. For the other 32% of trials, the cursor and cued bar were on opposite sides, with the cursor on the high probability side on 16% of the trials and on the low probability side on the other 16%. In the unattended condition, the direction of the arrow cued the expected side of the cursor on 20% of the trials. As such, the cursor and cued bar were on the same on only 32% of the trials, 16% of which on the high probability side and 16% on the low probability side. For the other 68% of trials, the cursor and cued bar were on opposite sides, with the cursor on the high probability side on 64% of the trials and on the low probability side on the other 4%.

Each participant completed two blocks of trials, one for the attended condition and one for the unattended condition. Each block included 10 baseline trials, 10 familiarization trials with clamped feedback, and 100 perturbation trials. The order of the attended and unattended blocks, the direction of the arrows, and the target positions (+/-45°, +/-135°) associated with attended/unattended session were counterbalanced across participants.

In Experiment 2b, two cursors were presented, one on each side of the target. The arrow pointed in one direction on 80% of the trials and in the other direction in 20% of the trials. Should there be any attention effect, it could accumulate across trials. In the control session, the direction of the cue was randomized (50/50 session). The order of the blocks, the direction of the arrow, and the target positions (+/-45°, +/- 135°) were counterbalanced across participants.

#### Experiment 3

Experiments 1 and 2 focused on the influence of spatial attention on implicit adaptation. In Experiment 3, we shift our focus, asking if adaptation is influenced by the availability of cognitive resources (Otsuka & Kawaguchi, 2007; Prull et al., 2016; Song, 2019). Specifically, we assume that the secondary bar discrimination task draws on cognitive resources. If adaptation also draws on the same resource pool, then adaptation might be attenuated by the inclusion of the secondary task.

We addressed this question in Experiment 3 by comparing adaptation under single and dual-task conditions. A 15° clamped cursor was presented on 2/3 of the trials (CW/CCW counterbalanced) and absent on 1/3 of the trials. We used a secondary task here, simply asking participants to report the side of the clamped cursor feedback or report that it was not presented. Depending on the accuracy of the response, the message “Good” or “Wrong” was presented for 700 ms after each response. In the single-task condition, the same conditions were presented but now the participant was instructed to ignore the feedback cursor (as in a “standard” clamp experiment). To roughly equate the motor requirement with that of the dual-task condition (Fig S2), the participant was required to press the “e” key after each trial, followed by the message “Good”.

#### Experiment 4

Experiment 3 contains a confound in that the focus of attention (and likely fixation) at the end of the trial differs between the single and dual-task conditions: For the former, the focus is on the target whereas for the latter, the focus is on the feedback cursor. In Experiment 4 we modified the dual-task such that the focus of attention should always remain on the target. Specifically, the target changed to yellow or red concurrently with the onset of the clamped cursor. After 50 ms, the target color changed back to blue and remained visible for another 90 ms. Participants in the dual-task condition reported the target color. In the single-task control condition, the same stimulus events occurred but the participants were instructed to ignore the color change.

#### Experiment 5

In Experiment 5, we further examined whether the processes involved in implicit adaptation are impacted when the availability of cognitive resources is manipulated. We again used an orientation discrimination task, presenting a red bar at the target location with the onset of the feedback cursor. Both were visible for only 50 ms, with the red bar then transformed into a cross for 90 ms. To manipulate resources, we varied the difficult of the secondary task (Hirst & Kalmar, 1987; Wickens, 2020), asking the participant to either report the orientation of the bar on the current trial (0-back condition) or report if the orientation on the current trial matched the orientation of the bar on the previous trial (1-back condition). We also included a single-task control condition in which the participant was told to ignore the red bar.

#### Experiment 6

As another test of the central resource idea, we used an auditory pitch discrimination task as the secondary task in Experiment 6. At movement onset, a series of three 40 ms pure tones were played, separated by 50 ms. The pitch of two of the tones were 261 Hz (middle C), and the other was 522 Hz (an octave higher). The participant indicated the ordinal position of the odd tone by choosing from three options using key press (1st/2ed/3rd). Performance on the tone task was indicated by a feedback message (Good/Wrong) presented for 700 ms after each response.

#### Experiment 7

Our final test of the resource was similar to Experiment 4 but involved a within-participant design. On 50% of the trials, the target changed from blue to orange in synchrony with the onset of the clamped feedback cursor, and then back to blue when the cursor disappeared for 90 ms. For the other 50% of the trials, the target color remained blue throughout the trial. The participant reported if the target had changed color (yes/no) by key press.

#### Experiment 8

The results of Experiments 1-7 suggested an alternative account of how attention might impact implicit adaptation: Namely, that attention might alter the context and create scenarios in which adaptation is attenuated due to a contextual change (Avraham et al., 2022; Heald et al., 2021, 2022; Shea & Morgan, 1979). In particular, when the focus of attention during action planning (i.e., target onset) is different from the focus at the time of feedback processing (i.e., clamp onset), different internal models are created and adaptation will be attenuated due to imperfect generalization (Haruno et al., 2003; Heald et al., 2022; Howard et al., 2013).

We tested this hypothesis in Experiment 8 by manipulating the consistency of the focus of attention within and between trials. A red bar, oriented vertically or horizontally, was embedded within the target at target onset and disappeared at movement onset. A second bar, either with the same or different orientation appeared within the target at feedback onset. The participants were always asked to report the orientation of the second bar; as such, the first bar was task irrelevant, but we assume is processed with the target and thus provides a contextual cue.

Two groups of participants were tested. For one group, the orientation of the second bar on trial N-1 was always the same as the orientation of the first bar of trial N. For the other group, the orientation of the second bar on trial N-1 was always different from the orientation of the first bar of trial N. We hypothesized that adaptation would be attenuated in the second group since the internal model retrieved at the onset of trial n is different from the internal model that had been updated on the previous trial. For both groups, the two bars within a trial can be either the same or different (counterbalanced). In this manner, the full design contains four conditions regarding the consistency of the bar orientation within and across trials (AA-> A, AB->B, AB-> A, AA-> B, Fig 4a).

### Analysis

We first assessed the accuracy of each participant on the secondary task in the dual-task conditions. If the accuracy level was below 70%, we excluded the participant’s data from all analyses (∼20%). Our reasoning here was that these participants might not have paid sufficient attention to the secondary task, a prerequisite for assessing the impact of attention on implicit adaptation.

For the analyses of adaptation, hand angle was defined as the angle between a line from the start location to the target and a line from the start location to the hand when the movement amplitude of the cursor had reached the target distance. Angular values were transformed such that positive angles correspond to values in which the hand direction is in the opposite direction of the clamped feedback relative to the target, the signature of adaptation. We excluded trials in which the movement duration was longer than 500 ms or the reach error was larger than 90°. Participant with more than 30% invalid movements were excluded from all subsequent analyses (∼5%, see table S1).

To analyze adaptation in the trial-by-trial designs, we calculated the change of hand angle after each trial. Specially, the learning expressed on trial n is calculated by subtracting the hand angle on trial n+1 from the hand angle on trial n. We used two methods to analyze adaptation in the block design of Exp 2. First, we applied a cluster-based simulation test on the learning curve, looking for epochs that showed a significant difference between the attended and unattended conditions (Avraham et al., 2021; T. Wang, Avraham, et al., 2024). Second, we used a composite measure, averaging the hand angle data across the entire perturbation phase.

T-tests and ANOVAs were used for between condition comparisons. For the t-tests, we report the Bayes factor BF10, to indicate the ratio of the likelihood of the alternative hypothesis (H1) over the null hypothesis (H0)(Schönbrodt & Wagenmakers, 2018; Stefan et al., 2019). A BF10 larger than 3 suggests moderate evidence supporting H1, while a BF10 smaller than 0.33 suggests moderate evidence supporting H0. We also report Cohen’s d as an estimate of effect size. Prior to conducting the t-test or ANOVAs. We confirmed that the data met assumptions concerning Gaussian distribution and homoscedasticity. These assumptions were violated in the accuracy data. Thus, we replaced the t-test with a Mann-Whitney U test when comparing accuracy.

## Author contributions

T.W., R.B.I. contributed to the conceptual development of this project. J.L. collected the data. T.W. analyzed the data, prepared the figures, and wrote the initial draft of the paper, with all the authors involved in the editing process.

## Acknowledgments

RBI is funded by the NIH (grants NS116883 and NS105839).

## Supplementary

**Fig S1.**
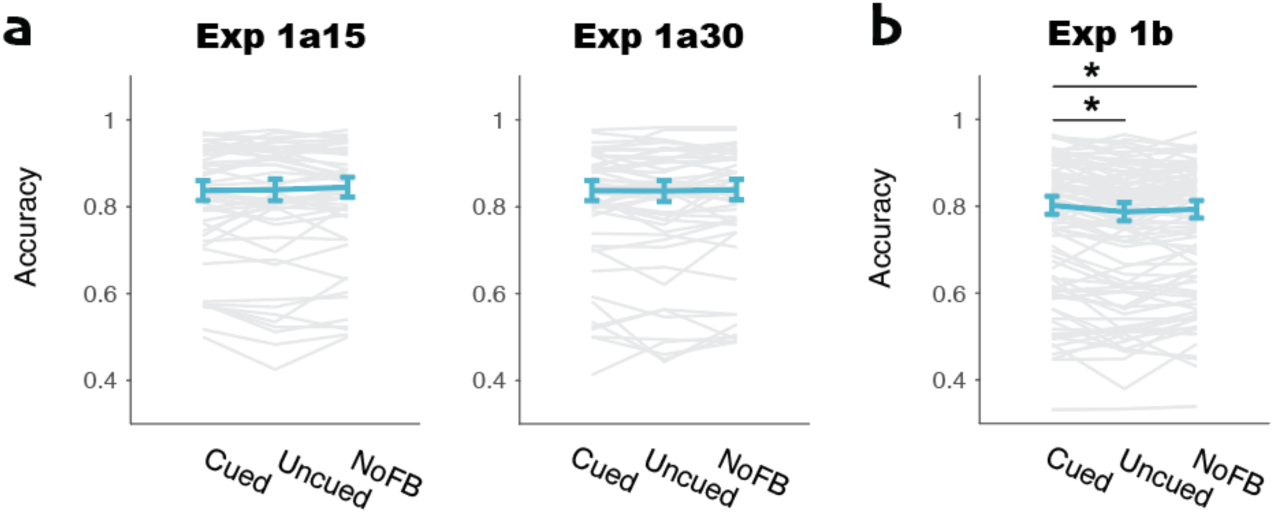
Second task performance of Experiment 1 for the attended, unattended, and no-feedback conditions. a) Accuracy on the bar discrimination task was similar across the three conditions in Experiment 1a, suggesting the feedback has minimal influence on the performance of the secondary task. b) In Experiment 1b, accuracy was higher in the attended condition compared to the other two conditions. This is likely because the red bar is presented on the white cursor in the attended condition, while the bar was presented on the black background in the other two conditions. The former is of higher visual contrast which might account for better discrimination performance in the attended condition. *,p<.05.

**Fig S2.**
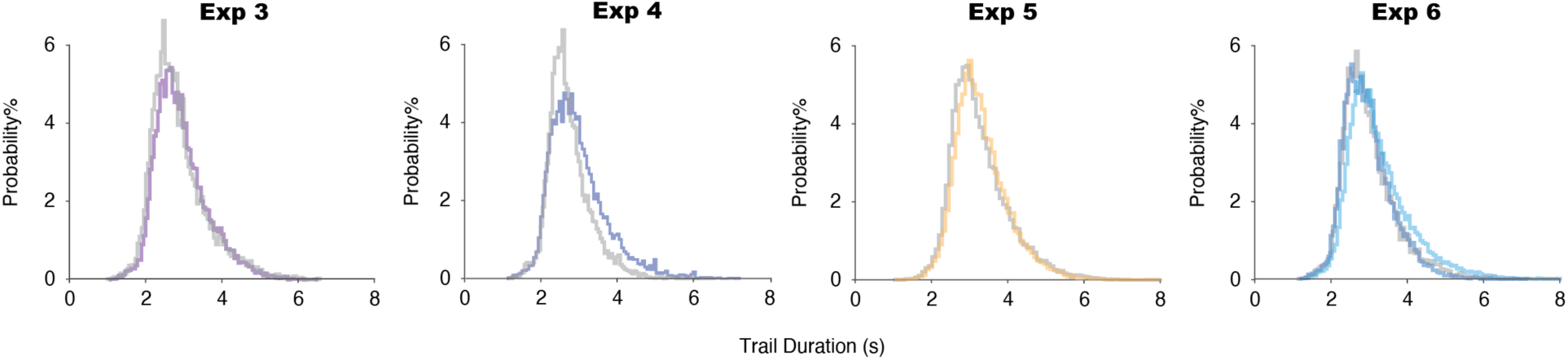
Distributions of trial duration were matched across the single and dual-task conditions in Experiments 3-6. Trial duration is defined as the interval between the onset of trial n and trial n+1.

**Fig S3.**
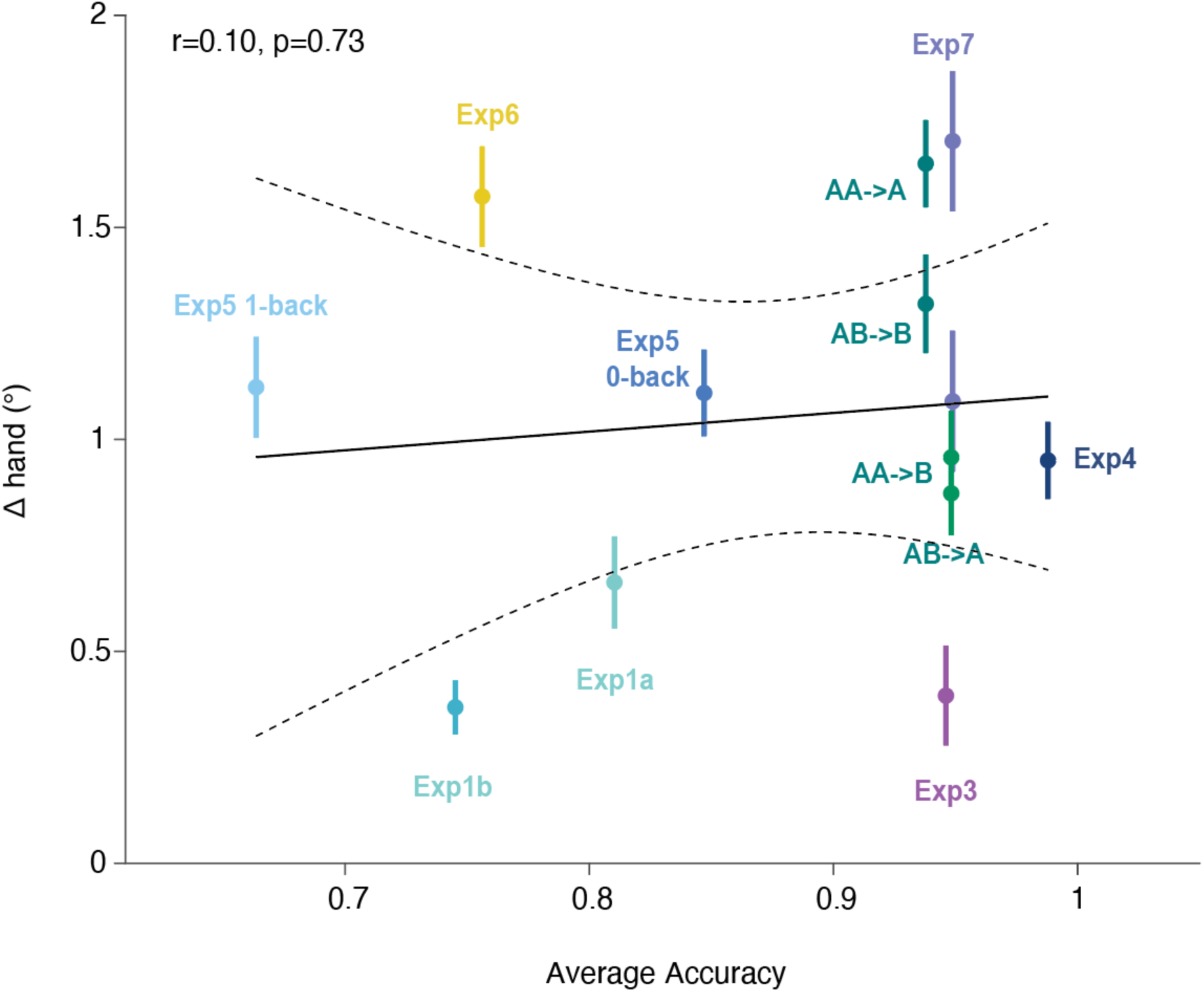
Post-hoc analysis of secondary task difficulty and magnitude of implicit adaptation. As our proxy of difficulty, we used secondary task accuracy. Δhand is plotted as a function of the accuracy. The black line indicates the best-fitted linear model, and the dash lines indicates the 95% confident interval of model. The data suggest minimal correlation between these two variables, providing further evidence that adaptation is not modulated by secondary task difficulty.

**Table S1.** Sample number of each Experiment.

| Condition | All Participants | Valid Participant |
| --- | --- | --- |
| Experiment 1a 15 | 50 | 35 |
| Experiment 1a 30 | 50 | 38 |
| Experiment 1b | 95 | 58 |
| Experiment 1c | 99 | 60 |
| Experiment 2a | 79 | 64 |
| Experiment 2b | 96 | 48 |
| Experiment 3 single | 34 | 32 |
| Experiment 3 dual | 34 | 32 |
| Experiment 4 single | 40 | 36 |
| Experiment 4 dual | 30 | 27 |
| Experiment 5 single | 33 | 32 |
| Experiment 5 0-back | 40 | 34 |
| Experiment 5 1-back | 56 | 20 |
| Experiment 6 single | 30 | 29 |
| Experiment 6 dual | 50 | 32 |
| Experiment 7 | 23 | 23 |
| Experiment 8 | 90 | 80 |
| All | 929 | 689 |

## Reference

1. Avraham, G., Morehead, J. R., Kim, H. E., & Ivry, R. B. (2021). Reexposure to a sensorimotor perturbation produces opposite effects on explicit and implicit learning processes. PLoS Biology, 19(3), e3001147.

2. Avraham, G., Taylor, J. A., Breska, A., Ivry, R. B., & McDougle, S. D. (2022). Contextual effects in sensorimotor adaptation adhere to associative learning rules. ELife, 11, e75801.

3. Bromberg, Z., Donchin, O., & Haar, S. (2019). Eye Movements during Visuomotor Adaptation Represent Only Part of the Explicit Learning. ENeuro, 6(6). 10.1523/ENEURO.0308-19.2019

4. Dunifon, C. M., Rivera, S., & Robinson, C. W. (2016). Auditory stimuli automatically grab attention: Evidence from eye tracking and attentional manipulations. Journal of Experimental Psychology. Human Perception and Performance, 42(12), 1947–1958.

5. Godden, D. R., & Baddeley, A. D. (1975). Context-dependent memory in two natural environments: On land and underwater. British Journal of Psychology (London, England: 1953), 66(3), 325–331.

6. Haruno, M., Wolpert, D. M., & Kawato, M. (2003). Hierarchical MOSAIC for movement generation. International Congress Series / Excerpta Medica, 1250, 575–590.

7. Heald, J. B., Lengyel, M., & Wolpert, D. M. (2021). Contextual inference underlies the learning of sensorimotor repertoires. Nature, 600(7889), 489–493.

8. Heald, J. B., Lengyel, M., & Wolpert, D. M. (2022). Contextual inference in learning and memory. Trends in Cognitive Sciences. 10.1016/j.tics.2022.10.004

9. Hirst, W., & Kalmar, D. (1987). Characterizing attentional resources. Journal of Experimental Psychology. General, 116(1), 68–81.

10. Howard, I. S., Wolpert, D. M., & Franklin, D. W. (2013). The effect of contextual cues on the encoding of motor memories. Journal of Neurophysiology, 109(10), 2632–2644.

11. Howard, I. S., Wolpert, D. M., & Franklin, D. W. (2015). The value of the follow-through derives from motor learning depending on future actions. Current Biology: CB, 25(3), 397–401.

12. Im, H. Y., Bédard, P., & Song, J.-H. (2016). Long lasting attentional-context dependent visuomotor memory. Journal of Experimental Psychology. Human Perception and Performance, 42(9), 1269– 1274.

13. Kasuga, S., Hirashima, M., & Nozaki, D. (2013). Simultaneous processing of information on multiple errors in visuomotor learning. PloS One, 8(8), e72741.

14. Kim, H. E., Avraham, G., & Ivry, R. B. (2021). The Psychology of Reaching: Action Selection, Movement Implementation, and Sensorimotor Learning. Annual Review of Psychology, 72(1), 61–95.

15. Kim, H. E., Morehead, J. R., Parvin, D. E., Moazzezi, R., & Ivry, R. B. (12/2018). Invariant errors reveal limitations in motor correction rather than constraints on error sensitivity. Communications Biology, 1(1), 19.

16. Kluzik, J., Diedrichsen, J., Shadmehr, R., & Bastian, A. J. (2008). Reach adaptation: what determines whether we learn an internal model of the tool or adapt the model of our arm? Journal of Neurophysiology, 100(3), 1455–1464.

17. Krakauer, J. W., Ghilardi, M. F., & Ghez, C. (1999). Independent learning of internal models for kinematic and dynamic control of reaching. Nature Neuroscience, 2(11), 1026–1031.

18. Krakauer, John W. (2009). Motor learning and consolidation: the case of visuomotor rotation. Advances in Experimental Medicine and Biology, 629, 405–421.

19. Krakauer, John W., Mazzoni, P., Ghazizadeh, A., Ravindran, R., & Shadmehr, R. (2006). Generalization of motor learning depends on the history of prior action. PLoS Biology, 4(10), e316.

20. Krakauer, John W., Pine, Z. M., Ghilardi, M.-F., & Ghez, C. (2000). Learning of Visuomotor Transformations for Vectorial Planning of Reaching Trajectories. The Journal of Neuroscience: The Official Journal of the Society for Neuroscience, 20(23), 8916–8924.

21. Liddy, J., & Song, J.-H. (2022). Implicit visuomotor adaptation is modulated by the attentional demands of a secondary task. Journal of Vision, 22(14), 4239–4239.

22. Logan, G. D. (1996). The CODE theory of visual attention: an integration of space-based and object-based attention. Psychological Review, 103(4), 603–649.

23. Mazzoni, P., & Krakauer, J. W. (2006). An implicit plan overrides an explicit strategy during visuomotor adaptation. The Journal of Neuroscience: The Official Journal of the Society for Neuroscience, 26(14), 3642–3645.

24. Morehead, J. R., Taylor, J. A., Parvin, D. E., & Ivry, R. B. (2017). Characteristics of Implicit Sensorimotor Adaptation Revealed by Task-irrelevant Clamped Feedback. Journal of Cognitive Neuroscience, 29(6), 1061–1074.

25. Ogasa, K., Yokoi, A., Okazawa, G., Nishigaki, M., Hirashima, M., & Hagura, N. (2024). Decision uncertainty as a context for motor memory. Nature Human Behaviour. 10.1038/s41562-024-01911-x

26. Otsuka, S., & Kawaguchi, J. (2007). Divided attention modulates semantic activation: evidence from a nonletter-level prime task. Memory & Cognition, 35(8), 2001–2011.

27. Pennycuick, C. (1996). Wingbeat frequency of birds in steady cruising flight: new data and improved predictions. The Journal of Experimental Biology, 199(Pt 7), 1613–1618.

28. Poh, E., Al-Fawakari, N., Tam, R., Taylor, J. A., & McDougle, S. D. (2021). *Generalization of motor learning in psychological space*. Neuroscience. http://biorxiv.org/lookup/doi/10.1101/2021.02.09.430542

29. Prull, M. W., Lawless, C., Marshall, H. M., & Sherman, A. T. K. (2016). Effects of divided attention at retrieval on conceptual implicit memory. Frontiers in Psychology, 7, 5.

30. Rajaram, S., Srinivas, K., & Travers, S. (2001). The effects of attention on perceptual implicit memory. Memory & Cognition, 29(7), 920–930.

31. Raymond, J. L., & Lisberger, S. G. (1998). Neural learning rules for the vestibulo-ocular reflex. The Journal of Neuroscience: The Official Journal of the Society for Neuroscience, 18(21), 9112–9129.

32. Redding, G. M., & Wallace, B. (1996). Adaptive spatial alignment and strategic perceptual-motor control. Journal of Experimental Psychology. Human Perception and Performance, 22(2), 379–394.

33. Roelfsema, P. R., Lamme, V. A., & Spekreijse, H. (1998). Object-based attention in the primary visual cortex of the macaque monkey. Nature, 395(6700), 376–381.

34. Schmitter-Edgecombe, M. (1996). The effects of divided attention on implicit and explicit memory performance. Journal of the International Neuropsychological Society: JINS, 2(2), 111–125.

35. Schönbrodt, F. D., & Wagenmakers, E.-J. (2018). Bayes factor design analysis: Planning for compelling evidence. Psychonomic Bulletin & Review, 25(1), 128–142.

36. Shadmehr, R., & Krakauer, J. W. (2008). A computational neuroanatomy for motor control. Experimental Brain Research. Experimentelle Hirnforschung. Experimentation Cerebrale, 185(3), 359–381.

37. Shea, J. B., & Morgan, R. L. (1979). Contextual interference effects on the acquisition, retention, and transfer of a motor skill. Journal of Experimental Psychology. Human Learning and Memory, 5(2), 179–187.

38. Smith, S. M. (1984). A comparison of two techniques for reducing context-dependent forgetting. Memory & Cognition, 12(5), 477–482.

39. Smith, S. M., & Vela, E. (2001). Environmental context-dependent memory: a review and meta-analysis. Psychonomic Bulletin & Review, 8(2), 203–220.

40. Song, J.-H. (2019). The role of attention in motor control and learning. Current Opinion in Psychology, 29, 261–265.

41. Song, J.-H., & Bédard, P. (2015). Paradoxical benefits of dual-task contexts for visuomotor memory. Psychological Science, 26(2), 148–158.

42. Stefan, A. M., Gronau, Q. F., Schönbrodt, F. D., & Wagenmakers, E.-J. (2019). A tutorial on Bayes Factor Design Analysis using an informed prior. Behavior Research Methods, 51(3), 1042–1058.

43. Taylor, J. A., Krakauer, J. W., & Ivry, R. B. (2014). Explicit and Implicit Contributions to Learning in a Sensorimotor Adaptation Task. Journal of Neuroscience, 34(8), 3023–3032.

44. Taylor, Jordan A., & Ivry, R. B. (2014). Cerebellar and prefrontal cortex contributions to adaptation, strategies, and reinforcement learning. Progress in Brain Research, 210, 217–253.

45. Taylor, Jordan A., & Thoroughman, K. A. (2007). Divided attention impairs human motor adaptation but not feedback control. Journal of Neurophysiology, 98(1), 317–326.

46. Thoroughman, K. A., Fine, M. S., & Taylor, J. A. (2007). Trial-by-trial motor adaptation: a window into elemental neural computation. Progress in Brain Research, 165, 373–382.

47. Tsay, J., Parvin, D. E., Dang, K. V., Stover, A. R., Ivry, R. B., & Morehead, J. R. (2024). Implicit adaptation is modulated by the relevance of feedback. Journal of Cognitive Neuroscience, 36(6), 1206–1220.

48. Tsay, J. S., Avraham, G., Kim, H. E., Parvin, D. E., Wang, Z., & Ivry, R. B. (2021). The effect of visual uncertainty on implicit motor adaptation. Journal of Neurophysiology, 125(1), 12–22.

49. Tsay, J. S., Ivry, R. B., Lee, A., & Avraham, G. (2021). Moving outside the lab: The viability of conducting sensorimotor learning studies online. *Neurons, Behavior*, Data Analysis, and Theory. 10.51628/001c.26985

50. Tsay, J. S., Parvin, D. E., & Ivry, R. B. (2020). Continuous reports of sensed hand position during sensorimotor adaptation. Journal of Neurophysiology, 124(4), 1122–1130.

51. Wang, J., & Sainburg, R. L. (2005). Adaptation to visuomotor rotations remaps movement vectors, not final positions. The Journal of Neuroscience: The Official Journal of the Society for Neuroscience, 25(16), 4024–4030.

52. Wang, T., Avraham, G., Tsay, J. S., Thummala, T., & Ivry, R. B. (2024). Advanced feedback enhances sensorimotor adaptation. Current Biology: CB. 10.1016/j.cub.2024.01.073

53. Wang, T., & Ivry, R. B. (2023). A Cerebellar Population Coding Model for Sensorimotor Learning. BioRxiv : The Preprint Server for Biology. 10.1101/2023.07.04.547720

54. Wang, T., Luo, Y., Ivry, R. B., Tsay, J. S., Pöppel, E., & Bao, Y. (2023). A unitary mechanism underlies adaptation to both local and global environmental statistics in time perception. PLoS Computational Biology, 19(5), e1011116.

55. Wang, T., Morehead, R., Tsay, J. S., & Ivry, R. B. (2024). The Origin of Movement Biases During Reaching. In bioRxiv (p. 2024.03.15.585272). 10.1101/2024.03.15.585272

56. Wang, T. S. L., Martinez, M., Festa, E. K., Heindel, W. C., & Song, J.-H. (2022). Age-related enhancement in visuomotor learning by a dual-task. Scientific Reports, 12(1), 5679.

57. Wickens, C. D. (2020). Processing resources and attention. In Multiple-task performance (pp. 3–34). CRC Press.

58. Wolters, G., & Prinsen, A. (1997). Full versus divided attention and implicit memory performance. Memory & Cognition, 25(6), 764–771.

59. Zhou, W., Fitzgerald, J., Colucci-Chang, K., Murthy, K. G., & Joiner, W. M. (2017). The temporal stability of visuomotor adaptation generalization. Journal of Neurophysiology, 118(4), 2435–2447.

